# K9HeartCircDB: A circRNA Atlas of Tachypacing-Induced Canine Dilated Cardiomyopathy

**DOI:** 10.64898/2026.06.16.732655

**Authors:** Chinmaya Chinmaya, Tanvi Sinha, Noemi Nisini, Tao Wang, Suriya Muthukumaran Natarajaseenivasan, Remus Berretta, Amit Kumar Rai, Amaresh Chandra Panda, John W. Elrod, Raj Kishore, Steven R. Houser, Fabio A. Recchia, Venkata Naga Srikanth Garikipati

## Abstract

Cardiovascular disease (CVD) remains a leading cause of death worldwide. Dilated cardiomyopathy (DCM), a major cause of heart failure (HF), exhibits ventricular dilation, impaired systolic/diastolic function, arrythmias, and adverse cardiac remodeling. While genetic causes of DCM have been extensively studied, non-genetic and acquired forms of DCM-like HF are less well characterized, especially with respect to non-coding RNA regulation.

Circular RNAs (circRNAs) are stable, covalently closed non-coding RNAs that regulate cellular function via sequestering miRNAs, RNA-binding proteins, or translation. Their role in canine HF that recapitulates features of non-genetic DCM remains largely unexplored.

To address this, we developed K9HeartCircDB (https://www.k9heartcircdb.com/), a publicly accessible database that catalogs circRNAs expressed in canine left ventricular (LV) tissues under tachypacing-induced HF, a model of non-genetic DCM-like disease, and healthy control conditions. The online interface enables users to query and explore circRNAs based on exon composition, predicted miRNA binding sites, protein-coding potential, siRNA targets, and primer design for experimental validation. By providing an integrated and user-friendly platform for canine heart circRNA exploration, K9HeartCircDB offers a valuable resource to facilitate mechanistic and advance translational studies on non-genetic DCM-like disease.

## INTRODUCTION

Cardiovascular disease (CVD) remains a leading cause of morbidity and mortality worldwide despite significant advances in diagnostic and therapeutic approaches^1^. Amongst CVDs, heart failure (HF) continues to impose a major clinical burden due to its progressive nature, complex pathophysiology, and limited capacity for complete recovery^1,2^. Dilated cardiomyopathy (DCM), a major cause of HF, is characterized by ventricular dilation and systolic and diastolic dysfunction and is associated with adverse cardiac remodeling, arrhythmia, and premature death^3–5^. Although DCM is linked to mutations in sarcomeric or desmosomal genes, it can also arise from non-genetic causes ^5–7^. These non-genetic DCM forms remain comparatively under characterized at the molecular level. While canonical signaling has been well characterized, the contribution of non-coding RNAs^8,9^ to the initiation and progression of non-genetic DCM remains incompletely understood.

Circular RNAs (circRNAs) represent a distinct class of endogenous non-coding RNAs generated through back-splicing of precursor transcripts, producing covalently closed RNA molecules that lack 5′ caps and 3′ poly(A) tails. This circular structure enables high stability and resistance to exonuclease-mediated degradation^10,11^. Accumulating evidence indicates that circRNAs participate in diverse biological processes through mechanisms such as miRNA sponging, interaction with RNA-binding proteins (RBPs), modulation of transcription and splicing, as well as in cap-independent translation^11–18^. In the CVDs, circRNAs have been implicated in cardiac hypertrophy^15,19^, myocardial infarction^11,13^, fibrosis^20,21^, endothelial dysfunction^22,23^, and HF^24^. However, their landscape in large-animal models of non-genetic DCM HF remains poorly defined.

Large-animal models closely recapitulate human cardiac physiology and disease progression compared to small animal systems^25,26^. Among these, the tachypacing induced canine HF recapitulates key features of non-genetic DCM and has served as an important translational model for studying ventricular remodeling in acquired forms of DCM ^27,28^. High-throughput RNA sequencing in such models offers an opportunity to identify disease-associated circRNAs^10^, which warrants exploring a new avenue for identification of novel therapeutic targets and biomarkers. However, the lack of a dedicated, searchable resource for canine cardiac circRNAs limits the accessibility and utility for further exploration in research. To address this gap, we developed K9HeartCircDB, a publicly accessible database for exploring circRNAs expressed in canine left ventricular (LV) tissues from tachypacing-induced HF and control hearts. This user-friendly platform enables exploration of circRNA features and supports hypothesis generation and translational studies linking canine models with human non-genetic DCM.

## MATERIALS AND METHODS

### Canine tachypacing-induced HF

Male mongrel dogs (1-2 years old, 22-25 kg; n=4/group) were chronically instrumented with a solid-state pressure transducer (E-430001-IMP-10, Stellar Implantable Transmitter, TSE systems) placed in the LV through the apex, a Doppler flow transducer (20 MHz pulsed Doppler velocimeter) positioned around the left circumflex coronary artery, and two pacing leads attached to the LV free wall epicardial surface, as previously described^29,30^. Catheters and wires were tunneled subcutaneously to the interscapular region and protected by custom-made jackets worn by the dogs. After 10 days of recovery from surgery, the pacing protocol was initiated to induce HF. Dogs underwent LV pacing using an external pacemaker at 210 bpm for 3 weeks, followed by an increased pacing rate of 240 bpm for an additional week **(Fig.1)**. This pacing protocol induces DCM and compensated HF during the first 3 weeks, progressing to decompensated HF between days 27 and 30, with LV end-diastolic pressure reaching ≥25 mmHg. All dogs were humanely euthanized on day 28 following acquisition of in vivo echocardiographic and hemodynamic data ^29,30^. Left ventricular tissue samples were snap-frozen in liquid nitrogen and stored at -80 °C until analysis. The protocol followed the National Institutes of Health guidelines for the care and use of laboratory animals and was approved by the Institutional Animal Care and Use Committee of Temple University^28,31^. The tachypacing model induces HF that recapitulates key features of non-genetic DCM. Throughout the study, we refer to this model as HF.

**Figure 1.**
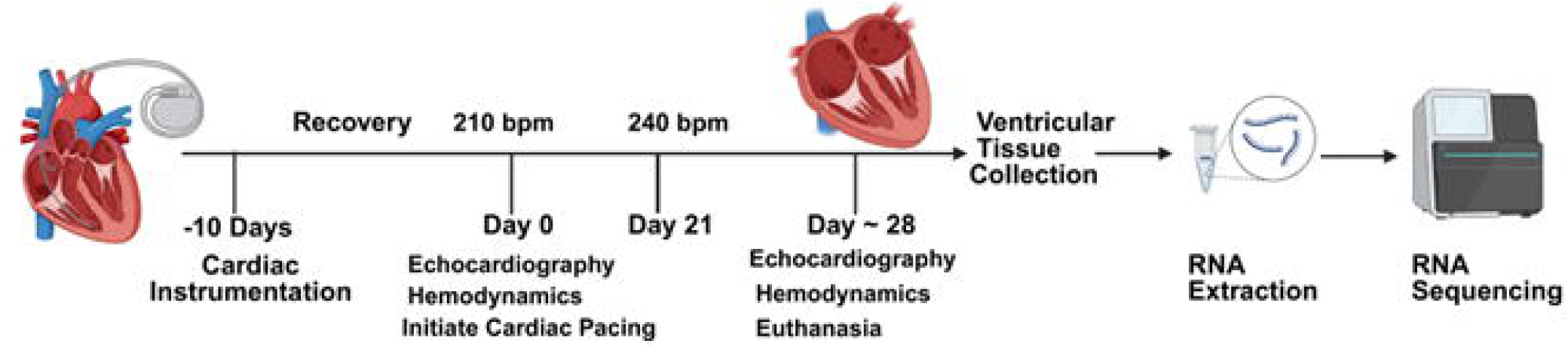
Study design and cardiac dysfunction in a tachypacing model of HF. Experimental protocol and study workflow. Cardiac instrumentation was performed at Day −10, followed by 28 days of rapid ventricular pacing (210–240 bpm) with interim echocardiographic assessment. At the study endpoint, ventricular tissue was collected for RNA extraction and sequencing.

### RNA isolation from the LV tissue

Total RNA (1□µg) was isolated from LV tissue using miRNeasy Mini Kit (Qiagen)^32,33^. For RNase R treatment, 1□μg of total RNA was incubated for 30□min at 37□°C with or without 2.5 U of RNase R (Epicentre Technologies, Madison, WI)^11^.

### Library Construction, Quality Control, and Sequencing

A total amount of 2 μg RNA per sample was used as an input material for the RNA sample preparations. Sequencing libraries were generated using RNA Library Prep Kit for Illumina following the manufacturer’s recommendations, and index codes were added to attribute sequences to each sample. Firstly, ribosomal RNA was removed, and rRNA free residue was cleaned up by ethanol precipitation. Subsequently, the linear RNA was digested with 3 U of RNase R per μg of RNA. Briefly, fragmentation was carried out using divalent cations under elevated temperature in First Strand Synthesis Reaction Buffer (5X). First strand cDNA was synthesized using random hexamer primer and M-MuLV Reverse Transcriptase (RNase H-). Second strand cDNA synthesis was subsequently performed using DNA Polymerase I and RNase H. In the reaction buffer, dNTPs with dTTP were replaced by dUTP. Remaining overhangs were converted into blunt ends via exonuclease/polymerase activities. After adenylation of the 3’ ends of DNA fragments, NEBNext Adaptor with hairpin loop structure was ligated to prepare for hybridization. To select cDNA fragments of preferentially 370-420 bp in length, the library fragments were purified with the AMPure XP system (Beverly, USA). Then 3 μL USER Enzyme was used with size-selected, adaptor-ligated cDNA at 37° C for 15 min, followed by 5 min at 95°C before PCR. Then, PCR was performed with Phusion High-Fidelity DNA polymerase, Universal PCR primers, and Index (X) Primer. Finally, products were purified, and library quality was assessed on the Agilent 5400 system and quantified by QPCR (1.5 nM). The Qualified libraries were pooled and sequenced on Illumina platforms with PE150 strategy in Novogene Bioinformatics Technology Co., Ltd (Beijing, China), according to effective library concentration and data amount required.

### CircRNA analysis

Quality of raw reads was assessed using the FastQC software (v 0.12.1), and adapters were trimmed using the fastp tool (v 0.23.2). Clean reads were mapped to the ROS Cfam 1.0 (GCF_014441545.1) reference genome using the STAR aligner (v 2.7.11b) with --chimSegmentMin 10 parameter. Circular RNAs were identified following the CIRCexplorer2 annotation pipeline (v 2.3.8). The mature spliced sequences of identified circRNAs were extracted using the BEDTools getfasta command-line (v 2.31.1). Differentially expressed circRNAs in LV heart-failure samples compared to the non-failure control were analyzed using the DESeq2 R package (v 1.44.0), with statistical significance kept at p-value less than 0.05 and log_2_FoldChange greater than 1.5.

### Divergent primer and siRNA design

Back-spliced junction sequence of circRNAs was used as a PCR template for designing divergent primers using the primer3 python script as described previously^34,35^. Depending on the spliced length of a given circRNA, the PCR template having size 200 nt or less was prepared by joining the 3′ and the 5′ halves of the mature circRNA sequence. Twenty-six nt back-spliced junction sequence of each circRNA was used as input to predict circRNA-specific siRNAs using the RNAxs perl package (http://rna.tbi.univie.ac.at/cgi-bin/RNAxs/), following a previously reported method ^34^.

### Cross-species conservation

The chromosomal coordinates of circRNAs curated in existing databases were retrieved and incorporated for cross-reference in the K9circRNA database^36–39^. Database IDs of the common dog circRNAs were identified by browsing the chromosomal coordinates available in public repositories, including circAtlas 2.0 and CIRCpedia3. The human (hg38) and mouse (mm10) circRNA ortholog IDs were identified by converting the Cfam 1.0 circRNA genomic coordinates into the respective species chromosomal locations using the UCSC liftOver tool^40^.

### Prediction of associated functional miRNAs

We downloaded the mature sequences of dog miRNAs from the miRBase database (accessed in July 2025), which was used to predict miRNA binding sites within the mature circRNA sequences using the miRanda (v 3.3a) tool with a score threshold of 150 or above ^36^.

### Potential protein-coding circRNAs

We utilized the CPAT (v 3.0.5) tool to predict the back-spliced junction spanning ORFs across the three-times concatenated mature sequences of LV failing and non-failing dog heart circRNAs (3XcircRNA). CPAT scores correlate directly to the protein-coding potential of circRNAs. We used IRESfinder (v 1.1.0) python package to predict the presence of internal ribosome entry sites (IRESs) in the 3XcircRNA sequences.

### Statistics

All statistical analyses were performed using GraphPad Prism. Data are presented as mean ± standard error of the mean (SEM). Comparisons between two paired groups were conducted using a two-sided paired Student’s t-test. A P value < 0.05 was considered statistically significant.

## RESULTS

### Cardiac Dysfunction and Pathological Remodeling in a Tachycardia-Induced HF Model

To confirm the development of HF, we performed echocardiographic and invasive hemodynamic assessments. At week 4, tachypaced animals exhibited an HF phenotype with DCM-like characteristics, including reduced LVEF and elevated LVEDP, consistent with progressive ventricular dysfunction and adverse cardiac remodeling, compared with controls **(Fig. 2A–B).**

**Figure. 2.**
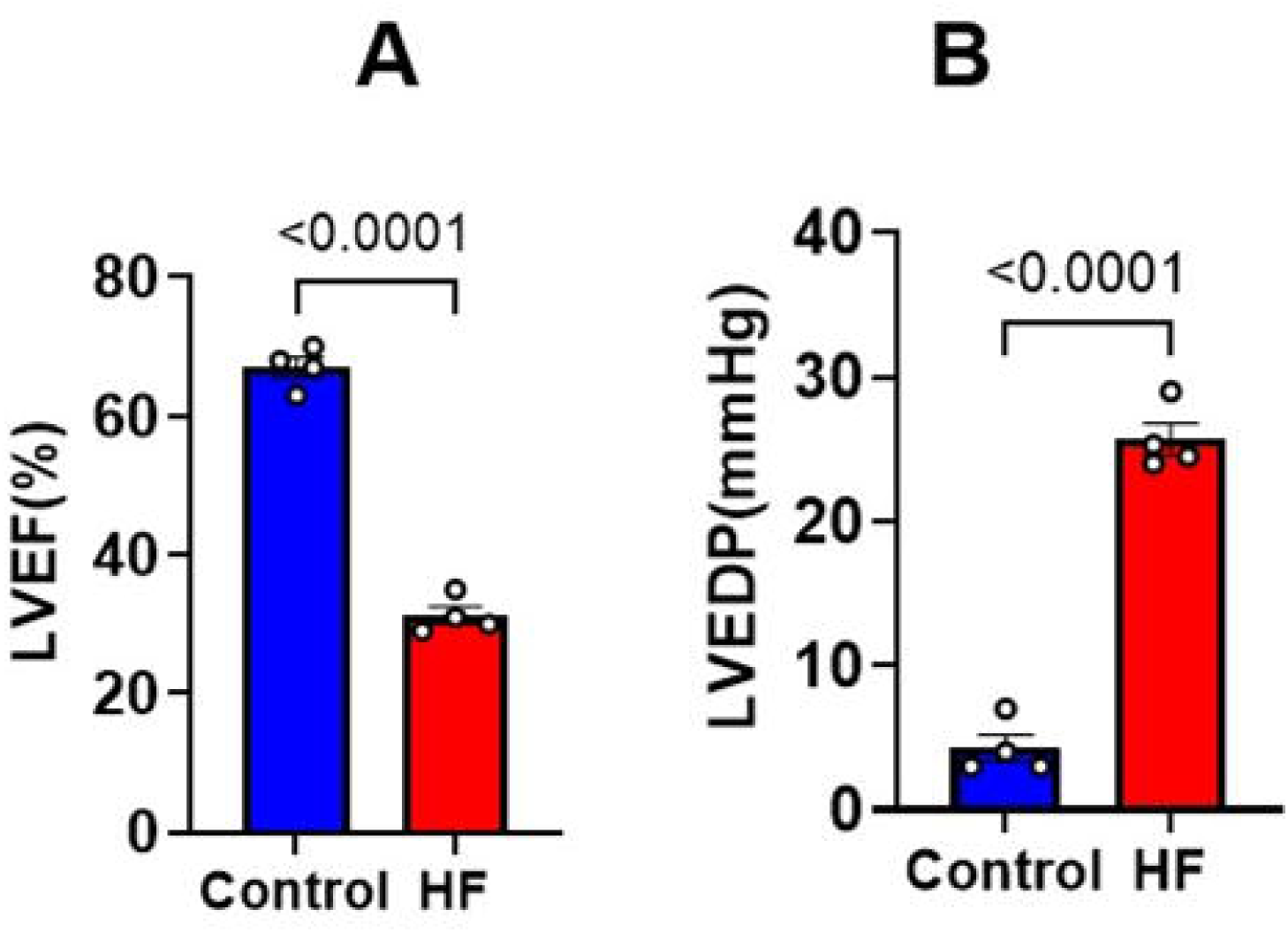
Cardiac functional and hemodynamic changes in control and HF groups. (A) Left ventricular ejection fraction (EF) and (B) left ventricular end-diastolic pressure (LVEDP) at week 4 in control and HF groups, demonstrating cardiac dysfunction and pathological remodeling (n = 4 per group).

### Characterization of circular RNAs and their properties in canine LV tissue from tachypacing-induced HF hearts

The RNA-seq datasets comprised quadruplicates of tachypacing induced heart-failure and control LV samples. A total of 37,667 unique circRNAs were identified in HF and control whole transcriptome datasets using the CIRCexplorer2 (v 2.3.8) pipeline **(Supplementary Table S1)**. Comparison between HF and control hearts yielded a list of 14,183 common circRNAs. Our database has identified 10,706 unique circRNAs in the HF group alongside 12,778 unique circRNAs in the control group. Most of the canine heart circRNAs were found to have a size smaller than 1 kb; exon count less than ten, and origin from the exonic regions of the genome. Differential expression profiling of the 14,183 common circRNAs was performed using the DESeq2 R package to find circRNAs with possible function in HF. A total of 50 circRNAs were identified with a p-value <0.05 and log_2_FC >1.5 that were significantly upregulated, and 52 circRNAs with a p-value <0.05 and log_2_FC >-1.5 that were significantly downregulated in HF compared to controls **(Fig.3 and Supplementary Table S2).** Furthermore, we conducted a cross-species conservation study of HF and control circRNAs identified in LV tissue using the genomic assembly conversion of dog circRNAs into corresponding human and mouse circRNAs with the UCSC LiftOver tool. It was found that a total of 12,724 circRNAs were conserved across all three species, providing an additional avenue for future multi-species conservation studies **(Supplementary Table S3).**

**Figure 3:**
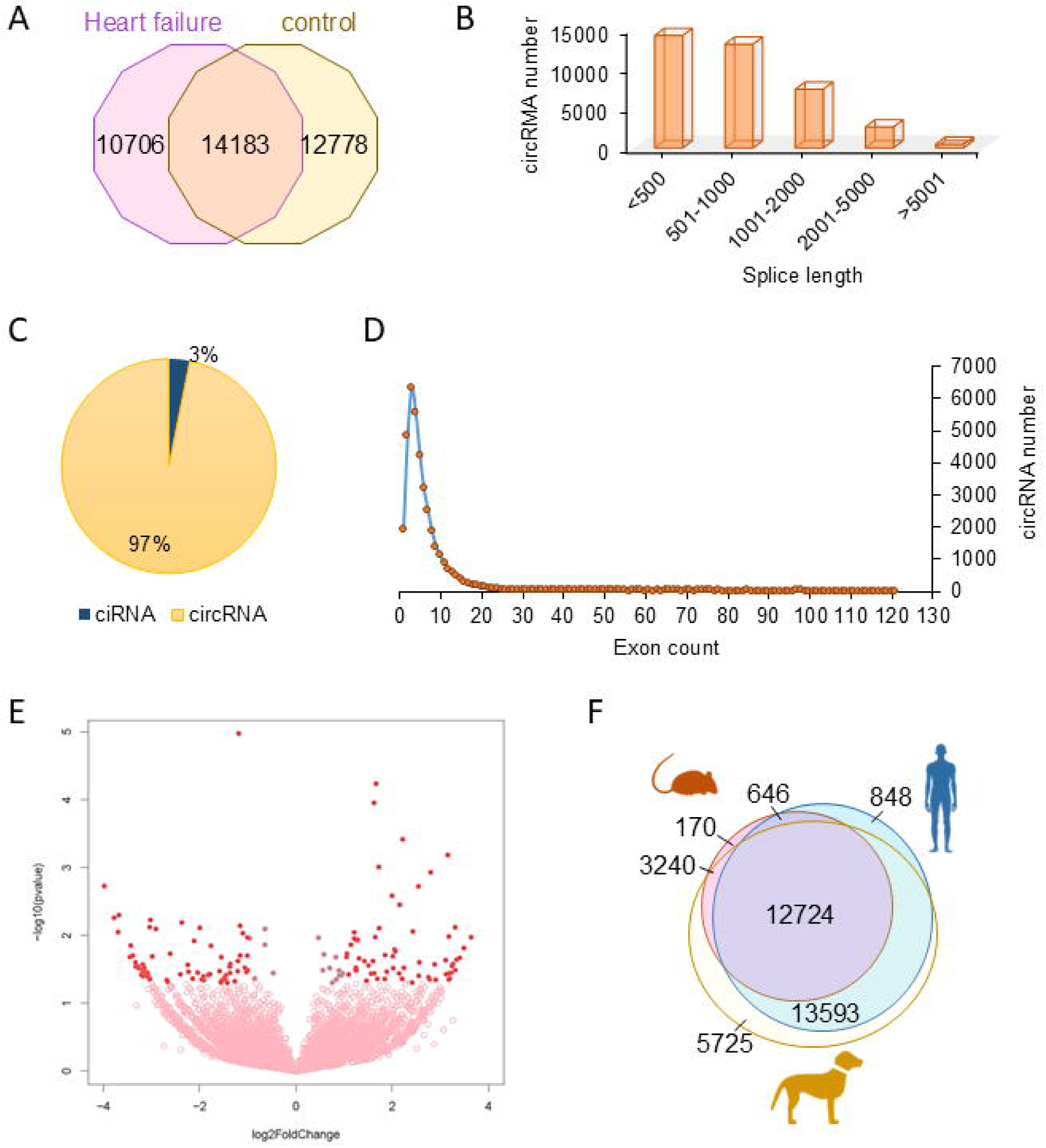
Identification and characterization of dog LV circRNAs in heart-failing and non-failing conditions: (A) Venn diagram showing common and unique circRNAs identified across the HF and NF groups. (B) Bar plot depicting the distribution of circRNAs based on spliced length range. (C) Pie chart showing percent of exonic (circ)RNAs and intronic (ci)RNAs. (D) Line plot showing circRNA population harboring one or more exons/introns. (E) Volcano plot showing distribution of differentially expressed circRNAs commonly identified in HF and NF conditions. (F) Venn diagram showing the number of cross-species conserved circRNAs identified in dog, human, and mouse.

### K9HeartCircDB search function and detailed information on each circRNA

We developed K9HeartCircDB, a comprehensive and user-friendly database that catalogs 37,667 circRNAs expressed in the LV tissue of failing and non-failing heart tissue. The platform allows the user to search for circRNA based on the circRNA ID and their chromosome coordinates listed in our database as ROS_Cfam_Chr_ID. User will be able to apply filters to refine results further. Instead of displaying all entries simultaneously, the interface enables selective expansion of matched entries, improving navigation and search **(Fig.4)**.

**Figure 4:**
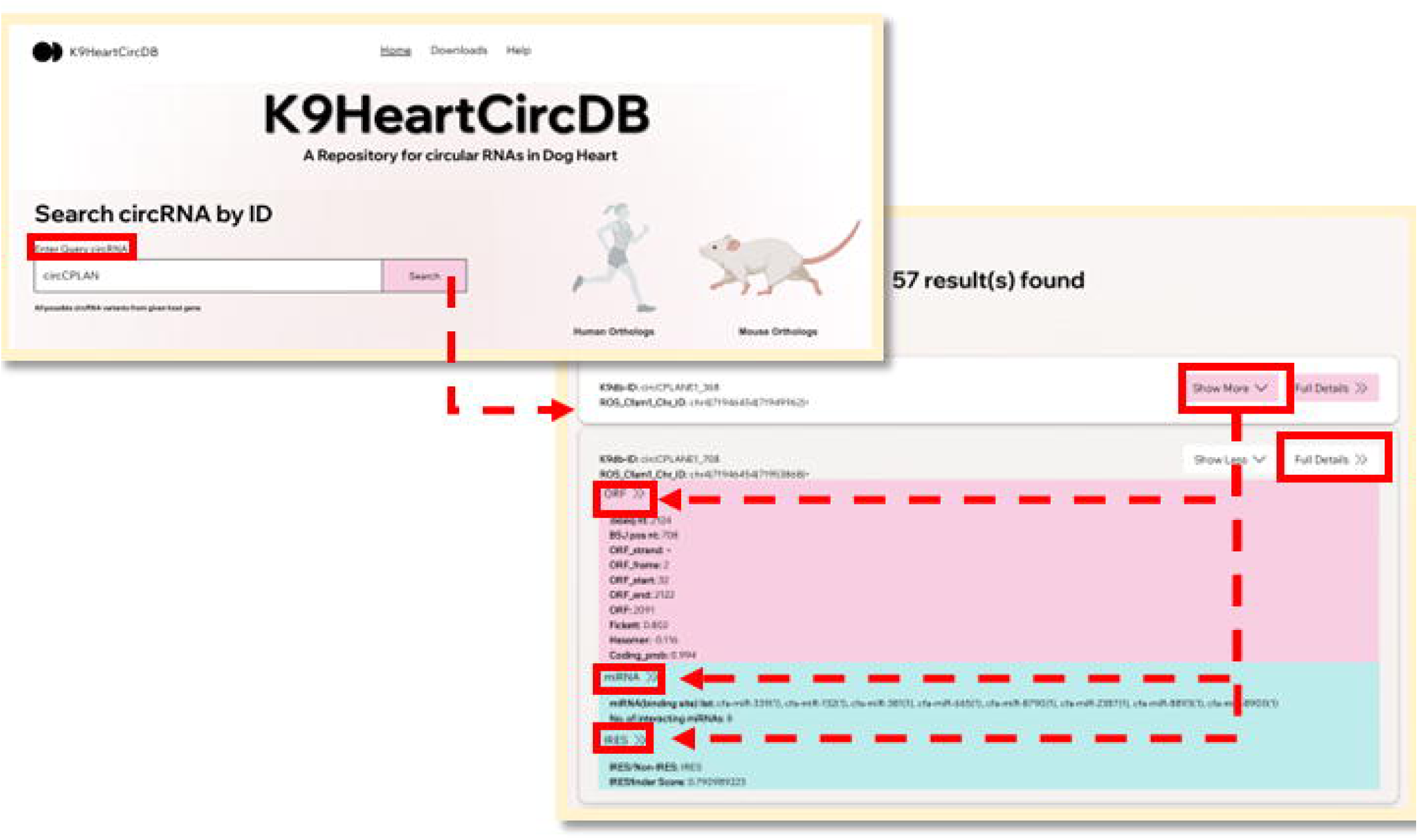
Overview of the K9HeartCircDB Web Interface. Users can query the database using a host gene symbol (e.g., “CPLANE”), a full circRNA identifier (e.g., “circCPLANE1”), or a prefix-based search (e.g., “circC”). The output displays all circRNAs associated with the queried gene or identifier.

The Results page provides information for each circRNA, including ORF, IRES-mediated translation sites, and miRNA binding sites. For information regarding primers and siRNAs, users can access “Full Details”, which represents K9CircDB-ID, divergent primers, interacting miRNAs, siRNAs, protein-coding potential, and PCR-related data across different databases. For example, a search for “circCPLANE” returns 56 circRNAs from CPLANE gene, such as *circCPLANE1_368, circCPLANE1_708, circCPLANE1_969* **(Fig.4)**

### Functional Annotation and Experimental Resource Integration in K9HeartCircDB

K9CircDB Database is a user-friendly and comprehensive resource that provides information on 37,667 circRNAs expressed in dog LV tissue in heart-failing and non-failing conditions. Besides the detailed circRNA data, K9CircDB offers characterization of circRNAs, including divergent primers, siRNAs, and related miRNAs sponged by the LV tissue circRNAs.

### Divergent primer for circRNA PCR

To validate circRNAs, K9CircDB provides information about at least one specific divergent primer pair for 37,588 circRNA K9dB_IDs, designed using the primer3.py script. The divergent primer specifically amplifies the target circRNA back-spliced junction sequence, which can be further validated by Sanger sequencing. Additionally, the database provides the circRNA mature sequence as well as the junction template sequence, which users can utilize to design divergent primers using Primer3 or the NCBI Primer-BLAST webserver **(Fig.5)**.

**Figure 5:**
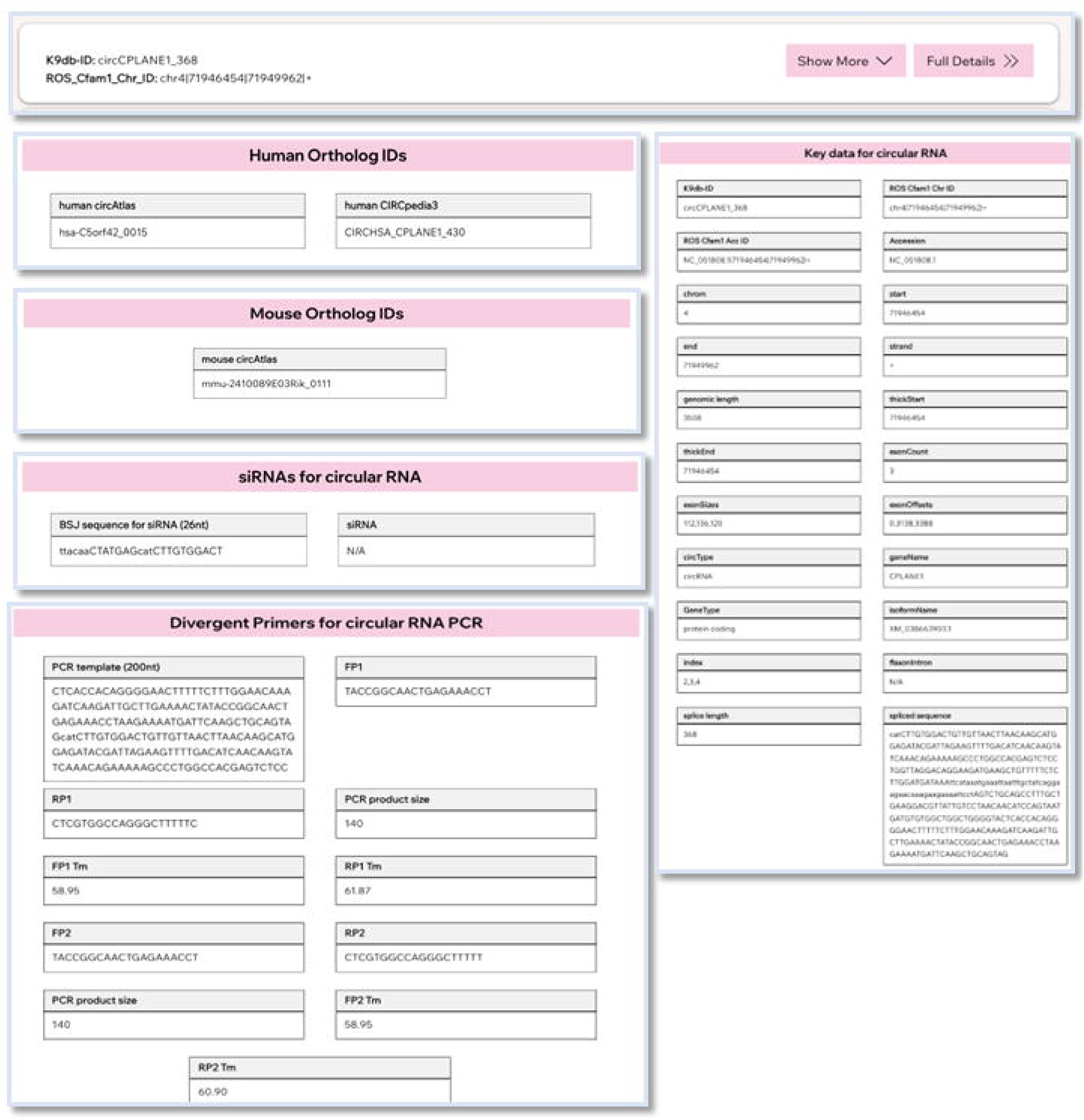
Interactive circRNA Annotation Panel Integrating Functional Features and Experimental Resources in K9HeartCircDB: Expanded circRNA Annotation Panel Displaying ORFs, miRNA Sites, IRES Elements, and Experimental Resources in K9HeartCircDB. Selecting “Show More” expands an entry to reveal a summary panel containing predicted open reading frame (ORF) sites, microRNA (miRNA) binding sites, and Internal Ribosome Entry Sites (IRES). Selecting “Full Details” opens a dedicated page providing comprehensive circRNA information, including genomic annotation, PCR primer sequences, siRNA targets, miRNA binding sites, protein-coding potential, and orthologous identifiers in human and mouse. Raw data files for ORF predictions, miRNA binding sites, and IRES elements are available for download for experimental use.

### siRNA for circRNA silencing

Rnaxs.pl is the tool that was utilized for the prediction of the siRNAs (http://rna.tbi.univie.ac.at/cgi-bin/RNAxs/). Out of the 37,667 circRNAs, our database hosts 25,842 K9dB_IDs with at least one predicted siRNA set, as shown in (**Fig.5)**. Out of the 25,842 identified, there were 3,770 circRNAs with a single siRNA set, 4,333 circRNAs with two siRNA sets, and 17,739 circRNAs with three siRNA sets. For example, three siRNA sequences targeting the back splice junction were predicted for *circUBL5_122*. Users are advised to confirm siRNA’s specificity using NCBI-BLAST. The siRNA sequences can be custom synthesized with two added nucleotides (dTdT) as 3′ DNA overhangs **(Fig.5)**

### Predicting functional miRNA targets

CircRNAs have been shown to sequester miRNAs, regulating a variety of cellular functions^16,41^. Using miRanda (v 3.3a), we identified 7,14,210 potential circRNA-miRNA interactions in LV circRNAs from failing and non-failing hearts, including 37,570 K9dB circRNA IDs. For example, circCPLANE1_708 harbors binding sites for different miRNAs, including *cfa-miR-339(1), cfa-miR-132(1), cfa-miR-381(1), cfa-miR-665(1), cfa-miR-8790(1), cfa-miR-2387(1), cfa-miR-8893(1), cfa-miR-8900(1),* where the number of binding sites per miRNA is shown enclosed in brackets.

The exact miRNA binding sites are displayed in **(Fig.5)** in a simplified filter format for quick access. This feature is incorporated into our database to reduce the time needed to sift through a list of miRNAs and only display the relevant ones.

### Protein Coding Potential of LV circRNAs

Although circRNAs lack the 5′ cap, some circRNAs can still be translated via IRES or m^6^A modifications. The Coding-Potential Assessment Tool (CPAT) analysis identified 17,554 circRNAs with at least one ORF spanning the back splice junction, suggesting that canine circRNAs could potentially translate into proteins. Additionally, we identified 27,602 dog LV circRNAs with potential IRES based on IRESfinder analysis^42^. For example, if the user input is NC_051809.1_24917649_24917727_PLUS or based on the gene name ciALKBH8_78, it is predicted to have a score of 0.984 according to IRESfinder analysis^39^ **(Fig.5).**

### UCSC track for circRNAs visualization

To visualize circular RNAs found in failing and non-failing heart conditions, we have added a feature allowing users to view them in the UCSC genome browser, accessible via the link provided on the database. The website also allows the user to navigate other databases to confirm the corresponding circRNA ID across different nomenclatures provided in other databases **(Fig.5).**

## DISCUSSION

The K9heartcircDB represents a comprehensive and uniquely curated resource comprising 37,667 circular RNAs identified in canine LV tissue under both HF and control conditions. By combining deep RNA sequencing with bioinformatic annotation, the database captures key features of circRNAs, including genomic coordinates, host genes, predicted microRNA (miRNA) interactions, and protein-coding potential. It also incorporates utility tools such as siRNA design and primer design information, making it useful not only for data exploration but also for experimental follow-up. In addition to cataloging novel circRNAs with consistent nomenclature, the database identifies those that are differentially expressed in the failing heart, providing a foundation for studying circRNA regulation in cardiac disease. Notably, the 26,317and 15,964 circRNAs exhibit homology to human and mouse counterparts, respectively, establishing a translational framework to study left ventricular dysfunction and HF, particularly in the setting of non-genetic DCM-like disease.

CircRNAs are known as regulators of gene expression in CVD, via sponging miRNA or RNA-binding proteins and in translation into peptides^11,13,14,43^. Although circRNA annotations have been described in human and rodent hearts^44,45^, their characterization in large-animal models has remained limited. Canine pacing-induced HF models closely resemble human features of non-genetic DCM. In this context, the **K9HeartCircDB** offers an opportunity to examine circRNAs in a model that more closely reflects human cardiac physiology and disease progression.

Earlier studies have reported circRNA expression in canine atrial tissue^46^, human dilated cardiomyopathy (DCM)^47,48^, and in circulation^49^. Although existing circular RNA databases^45,50^ provide annotated cardiac circRNAs in rodents and humans; their systematic characterization in larger animal particularly in the context of canine non-genetic DCM, remain limited. This gap is significant as small animal models do not completely recapitulate the molecular complexity in human cardiac disease. In this context, the canine tachypacing-induced HF model closely mirrors important features of human non-genetic DCM, including ventricular dilation, contractile dysfunction, and progressive remodeling. By using LV tissue of pacing-induced canine HF models, circRNA changes in the primary sites of contractile dysfunction, reflecting transcriptional changes to mechanical stress and remodeling. Furthermore, the identification of circRNAs conserved across species enables cross-species comparisons and functional interrogation in complementary model systems. Overall, this resource expands the current landscape of cardiac circRNAs in a large-animal model and provides a foundation for mechanistic and translational studies of left ventricular dysfunction by enabling systematic exploration of circRNA features in a platform that bridges canine models with human non-genetic DCM.

## LIMITATIONS

This database has several limitations. First, it is derived from bulk LV tissue, which may obscure cell–type–specific circRNA expression and limit resolution of cell-specific regulatory mechanisms. Second, circRNA identification and functional annotation, including predicted microRNA interactions and coding potential, are primarily based on *in silico* analyses and therefore require experimental validation. Future studies will be needed to confirm circRNA expression and functional relevance using targeted approaches. In addition, integration of cross-species comparisons with human and murine cardiac circRNA datasets will further enhance its translational relevance.

## Data availability statement

The datasets used in this study are provided on the database.

## Author contributions

## Acknowledgements

C.C.: Conception and design, C.C., T.S, N.N., S.N, R.B, T.W, A.K.R.: Collection and data analysis; A.K.R, A.C.P, J.W.E., R.K., F.A.R., V.N.S.G.: conception and design, final editing and approval of manuscript.

